# A high resolution view of adaptive events

**DOI:** 10.1101/429175

**Authors:** Han Mei, Barbara Arbeithuber, Marzia A. Cremona, Michael DeGiorgio, Anton Nekrutenko

## Abstract

Coadaptation between bacterial hosts and plasmids often involves a small number of highly reproducible mutations. Yet little is known about the underlying complex dynamics that leads to such a single “correct” solution. Observing mutations in fine detail along the adaptation trajectory is necessary for understanding this phenomenon. We studied coadaptation between *E. coli* and a common artificial plasmid, pBR322, in a continuous turbidostat culture. To obtain a high resolution picture of early adaptive events, we used a highly sensitive duplex sequencing strategy to directly observe and track mutations with frequencies undetectable with conventional methods. The observed highly reproducible trajectories are governed by clonal interference and show rapid increases in the frequencies of beneficial mutations controlling plasmid replication followed by a profound diversity crash corresponding to the emergence of chromosomal variants. To the best of our knowledge our study represents the first comprehensive assessment of adaptive processes at a very fine level of resolution. Our work highlights the hidden complexity of coadaptation, and provides an experimental and theoretical foundation for future studies.

## Introduction

Mutations continuously feed prokaryotic genetic diversity, which, when exploited by selection, enables microbes to colonize most environments, including our own bodies, and to defy antibiotic agents and host defences. Our understanding of these adaptive processes has been greatly advanced by experimental studies in bacteria^1–4^, bacteriophages ^5–7^, yeast ^8–10^, and other systems (for review see ^11–13^). The fate of beneficial mutations in clonal asexual populations depends on their supply (a product of population size and the mutation rate) and the distribution of fitness effects. If the supply of beneficial mutations is low relative to the selection strength, then a new variant quickly spreads and fixes. On the other hand, if mutational supply is high, then multiple coexisting variants interfere with each other—a phenomenon initially termed *periodic selection*^14^ and known today as clonal interference ^15–17^. Because population sizes of clonal asexual organisms are typically very large, there is a sufficient supply of beneficial mutations so that their frequencies are governed by clonal interference. Thus understanding the fine dynamics of clonal interference is necessary to predict adaptive events in such organisms^18,19^.

To gain this understanding, one would need to try tracking all mutations within a population as soon as they arise. This may be technically challenging. The emergence of modern sequencing technologies has created an opportunity to explore these underlying dynamics^2,4,7^. However, these techniques only allow the observation of evolutionary trajectories of mutations that have risen to frequencies of ~1%—a threshold at which the noise of Illumina sequencing (the most accurate of the currently available sequencing technologies) obscures the signal^20,21^. Such a threshold is too high to obtain reliable insight into the underlying genetic variability within a large population. For example, Good et al.^2^ used one milliliter of the overnight *E. coli* culture for sequencing time points from the long term evolution experiment. This volume conservatively contains ~10^8^ cells. As a result, at sequencing resolution of 1%, one will miss all sequence variants present in fewer than ~10^6^ cells. A profound gain in resolution has been made by the application of molecular barcodes within an experimental evolution setting^9,10,22^. These studies have shown that at the beginning of an experiment, a large pool of lineages carrying beneficial mutations is present within the frequency band between 10^−8^ and 10^−4^, and a vast majority of adaptive lineages have frequencies under 0.01% (well below what is detectable with conventional Illumina sequencing). These data provided us with invaluable insight into early adaptive events. However, they do not convey the nature of genetic variants, as the barcode counts are only proxies for the number of cells carrying a particular, yet unknown, beneficial mutation. Venkataram et al.^22^ recognized this limitation and used barcoding to select several hundred lineages carrying beneficial mutations for subsequent whole genome sequencing. Still, this is only the tip of the iceberg, with the majority of variants remaining below the surface.

A number of techniques have been recently developed to allow sequencing-based detection of very rare genetic changes that may be suitable to directly observe early adaptive events. Of these, duplex sequencing is the most sensitive, with a theoretical resolution threshold of <10^−7, 21,23^. It is based on using unique sequence tags to label individual molecules of the input DNA prior to preparation of Illumina sequencing libraries. During the amplification steps of library preparation, each of these molecules gives rise to multiple descendants. After sequencing, the descendants of each original DNA fragment are identified and grouped together using tags—i.e., one simply sorts tags in sequencing reads lexicographically, and all reads containing the same tag are bundled into *families*. These families (usually with at least three members) form single strand consensus sequences (SSCS) for the forward or the reverse strand, respectively. Complementary SSCS are then grouped to produce duplex consensus sequences (DCS). A true sequence variant is present in all reads within a family forming a duplex. In contrast, sequencing and amplification errors will manifest themselves as “polymorphisms” within a family, and so they can be identified and reliably removed. The resolution of duplex sequencing combined with the power of experimental evolution approaches provides a unique opportunity to directly observe adaptive events as they appear.

Observing emerging mutations and tracking their evolutionary trajectories requires an experimental system that satisfies two criteria. First, it must have a sufficiently small genome to allow for ultra-deep duplex sequencing. This is because the great dynamic range of duplex sequencing requires high depth (i.e., identifying variants with frequency of 10^−6^ requires sequencing depth of >10^−6^). Second, such a system needs to evolve on a timescale that fits into a short term experimental evolution setup, such as the one provided by chemostat or turbidostat devices. To satisfy these requirements, we selected one of the logistically simplest and exceptionally well understood systems–plasmid vector pBR322. It has a small genome of 4,361 bp, which is amenable to deep sequencing. As a synthetic plasmid it has no evolutionary history of being associated with *E. coli*. Transforming it into naïve *E. coli* cells triggers a series of adaptive events in the plasmid itself as well as in the host genome ^24–28^. These changes take place within as few as several hundred generations, which is an easily achievable timeframe for an experimental evolution experiment. Because acquisition of the plasmid incurs a metabolic burden on the host cells, it reduces growth rate relative to their plasmid-free isogenic counterparts^27^ and results in the rapid decline of transformed cells in antibiotic-free chemostat cultures^29^. pBR322 contains two antibiotic resistance genes encoding a (β-lactamase and aTetC efflux pump, conferring resistance to ampicillin and tetracycline, respectively. Incubating transformants in the presence of these antibiotics eliminates plasmid loss as these genes become essential for host survival. Furthermore, the plasmid-encoded *tetC* gene plays a critical role in the host-plasmid co-adaptation. Propagation of naïve cells transformed with *tetC*-containing plasmid changes the host phenotype to a state where the plasmid carriage is no longer detrimental, and this change occurs in only 500 generations^24^. The evolved host has higher fitness in the presence of the plasmid, and this effect, caused by mutation(s) on the bacterial chromosome, requires a plasmid carrying *tetC* such as pBR322. Biochemically, this may be explained by *tetC* assuming other roles beyond tetracycline efflux (e.g., potassium uptake^28^).

The coevolution between a host and a plasmid provides a well defined experimental setting for direct observation of clonal interference. When a population (large enough to allow for clonal interference to drive adaptation) of naïve cells transformed with the plasmid is propagated within an antibiotic-containing medium, a number of competing beneficial mutations arise and follow the dynamics that was observed by Levy et al.^9^ with multiple adaptive lineages co-existing at relatively low frequencies. Subsequently large effect mutation(s) sweep to fixation and lead to the crash of genetic variability within the population. Lenski et al. established that this large fitness effect change arises within ~500 generation^24^. Thus, by taking regular samples across the duration of the adaptation experiment, we were hoping to reconstruct the trajectory of adaptive events.

In this study we directly monitored the fate of beneficial mutations arising during coadaptation of a bacterial host and a plasmid in a continuous culture. Our objective was to test repeatability of early adaptation, to develop a simple deterministic model that would capture the observed behavior, and to generate predictions for future experiments.

## Results and Discussion

### Experimental evolution setup

We chose to use a turbidostat, a device that maintains constant cell density with no restrictions on the supply of nutrients, for this experimental evolution experiment, as it puts additional selective pressure on host cells to increase their maximum growth rate^30,31^. We selected *E. coli* DH_5_α F-*recA1* as the host, in which recombination is inhibited and conjugation is disabled. The cells were transformed with pBR322—a low copy number plasmid (15–20 per cell) ^32,33^ that minimizes the selection of traits favoring competition among the plasmid population^34^ and that carries two antibiotic resistance loci. The turbidostat was set up in the following way. At time point zero, the turbidostat was inoculated with an aliquot of overnight DH5α culture transformed with pBR322. To establish the time course of the adaptation, approximately ⅔ of the volume (~8 mL) was taken from the incubation vessel every 12 hours, constituting time points 1,2, 3, and so on. Plasmid DNA was isolated from that volume and subjected to duplex sequencing without any additional manipulation. Sampling such a relatively large volume allowed us to perform thorough population level sampling. Two experiments including two replicates each were performed: a short term (60 hours; replicates R1 and R2) and a long term (318 hours; replicates R6 and R7). Fig. 1 shows a graphical representation of our sampling strategy with the optical density at 600 nm (OD_600_) dips corresponding to sampling points. Duplex sequencing was performed at 0,12, 24, 36, 48, and 60 hour points in the short term experiments and at 72, 84,108,156, 240, and 318 hour points in the long term experiments. In each case, both replicates were sequenced for a total of 24 duplex sequencing datasets. In addition, we performed conventional high depth sequencing of the host genome before (naïve) and after (evolved) the long term experiments.

### Emergence of polymorphisms followed by variation crash

Frequencies and positions of single nucleotide variants (SNVs) and insertions/deletions (indels) called from duplex sequencing are shown in Table S1 and Fig. 2. The initial, monomorphic, samples (time point zero, short term experiments, replicates R1 and R2) contained five SNVs supported by a single duplex family each (implying that our sample contained only a single molecule carrying each SNV). These sites did not propagate through the experiment and disappear in subsequent time points. We call such positions *flickering* sites, which likely represent neutral variants lost due to genetic drift. Flickering sites are present in most (but not all) time points, and the vast majority of them are supported by only one duplex family. The other group of variants is represented by likely adaptive SNVs that persist through the time points. None of them appear in the initial sample, and their frequencies increase initially (short term experiment) and start to drop off as the experiment progresses (long term experiment; Fig. 2) so that, at the terminal time point of one of the replicates of the long term experiment, pBR322 reverses completely to the wild-type. These SNVs cluster exclusively in two regions within the replication origin of pBR322: positions 3,027-3,035 and 3,118. In total, all experiments yielded 145 SNVs. The mutational spectrum (Fig. S1) showed a transition-to-transversion ratio of 1:1.9, which matches that of the long term evolution experiment of Wielgoss et al.^35^. One insertion and two deletions were identified in total (Table S1). The insertion was located at the replication origin, and was observed only in replicate R7 of the long experiment. One deletion that was located outside the origin was found in multiple turbidostat runs. The other deletion was within the origin, and was only observed in the R_1_ replicate of the short experiment. Indels did not exhibit any frequency fluctuation, which could be due to the rate of indel mutations being generally lower than that of base-pair substitutions ^36–38^.

**Figure 1.**
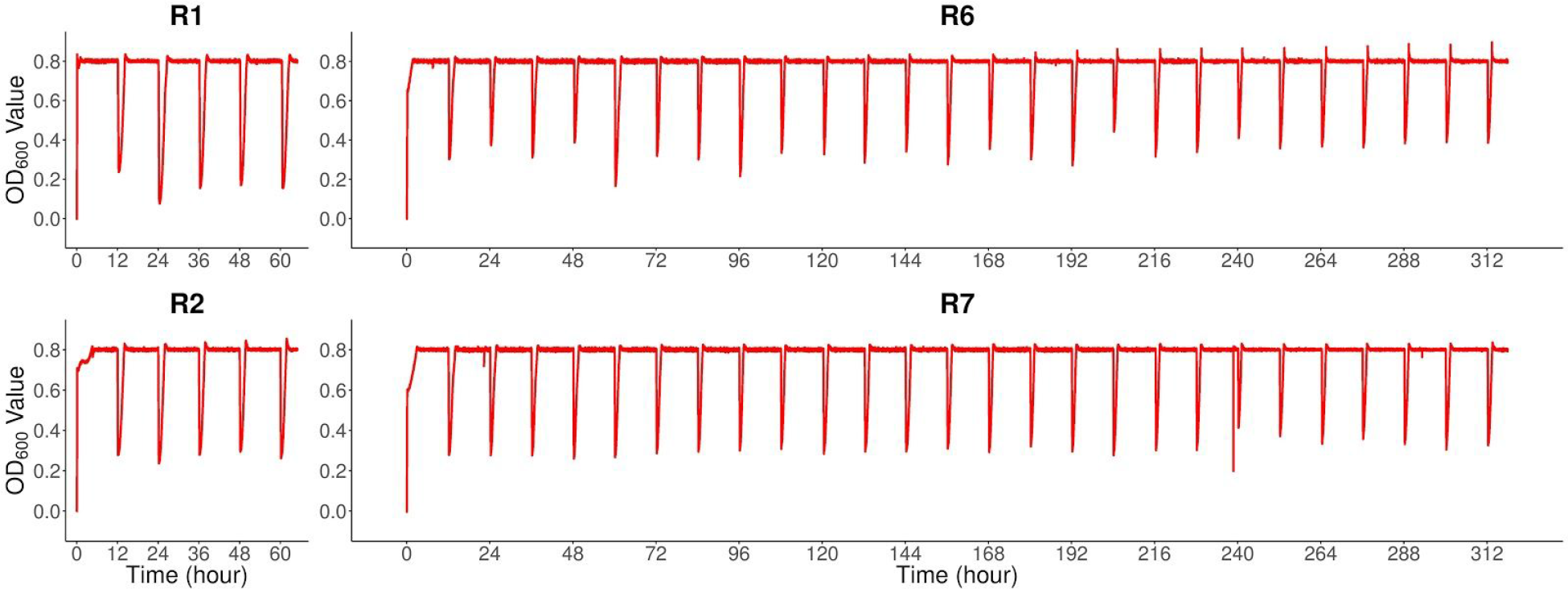
The four replicates of the short term (R1, R2) and long term (R6, R7) turbidostat runs. OD_600_ values were constantly monitored and maintained at 0.8. Samples were taken every 12 h. Dips in OD values represent sampling points. At each sampling point, ⅔ of the turbidostat volume was extracted for duplex sequencing.

**Figure 2.**
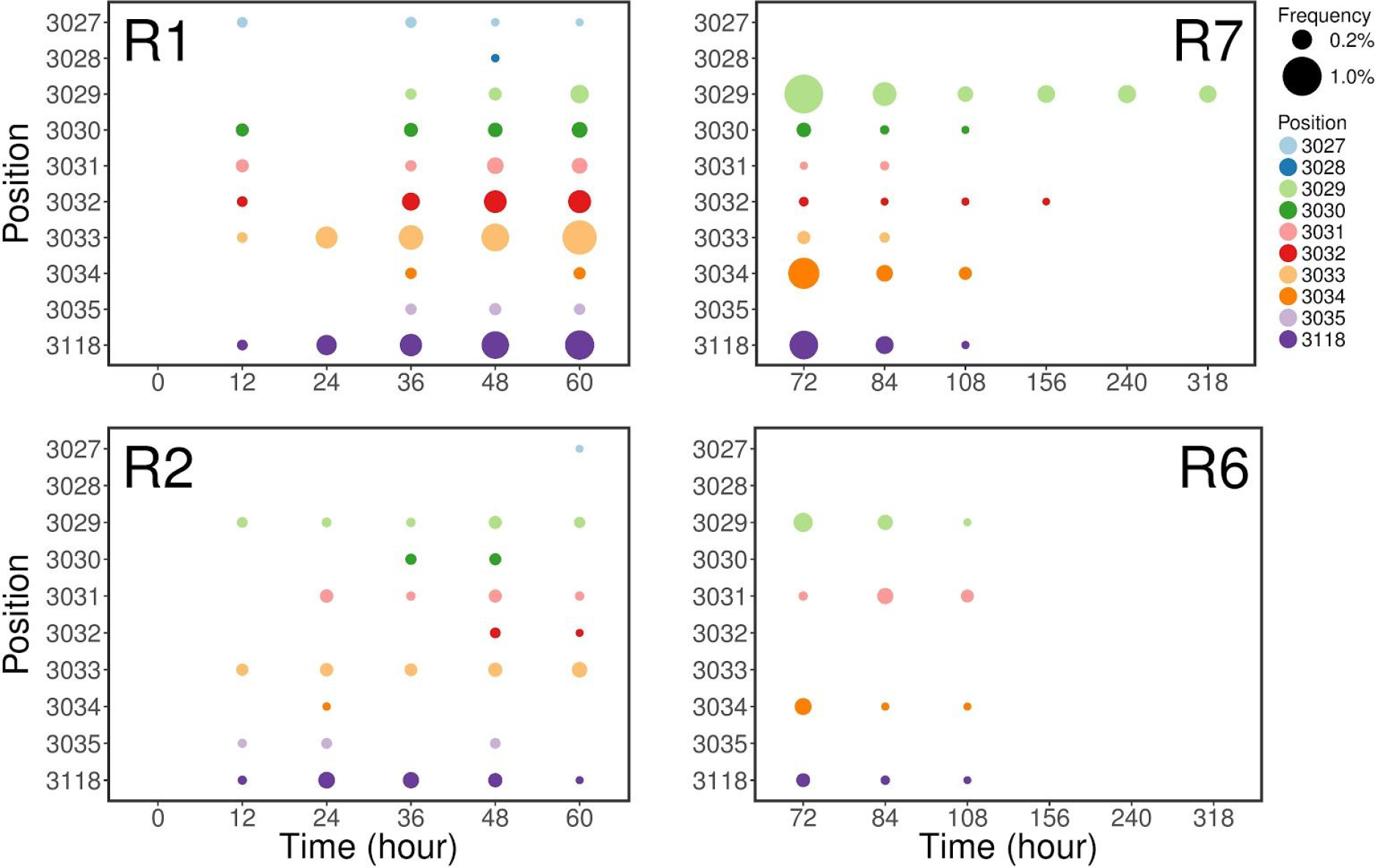
Locations and frequencies of variants detected at bases 3,027-3,035 and 3,118 on pBR322 with duplex sequencing. R1 and R2 denote the two replicates on short turbidostat run. R6 and R7 denote the two replicates of long turbidostat run. These variants contributed to change in plasmid copy number. Positions are projected to the y-axis, and colored individually. Sizes of the closed circles are proportional to the allele frequencies of corresponding variants.

Because the distance between SNVs located within the replication origin is shorter than the length of duplex reads, we examined our data for evidence of linkage. No linkage between any SNVs was observed. This is not unexpected because given the *E. coli* mutation rate is ~10^−10^−10^−9^ per nucleotide per generation^35–38^ and the highest frequency attained by these SNVs is ~0.9% it is highly unlikely to have two mutation events within a single molecule.

### Variable sites control plasmid copy number

The mechanism controlling pBR322 replication has been well studied^39^. Two complementary RNA molecules are encoded within the *ori* region: RNAI and RNAII. RNAII is first transcribed and hybridizes with plasmid DNA to form a primer complex for DNA replication. The other molecule—RNAI—is transcribed from the complementary strand. RNAI inhibits the formation of RNAII-DNA complexes, a replication primer, by interacting with RNAII via three loop regions. Substitutions within region 3,27-3,035 (Fig. S2A) were reported to increase pBR322 copy number^40^. This interval represents loop II’ of RNAI. Meanwhile, base 3,118 corresponds to the −35 promoter region of RNAII (Fig. S2B). Changes at this base were reported to increase the copy number as well^41^. The absolute majority of all variants found in this study are confined to these regions.

### Evolved cells have higher fitness conferred by chromosomal mutations

At each collection point we sampled ⅔ of volume from the incubation vessel, which was immediately topped-off with sterile medium diluting the remaining cells and causing optical density dips in Fig. 1. In each case, OD_600_ values quickly returned to the 0.8 threshold. The speed of this recovery can be used as a relative fitness estimate: the steeper the slope, the fitter the cells. This approach demonstrated consistent fitness increases over the course of turbidostat runs (Figs. S3 and S4, left panels). Contrasting the fitness of terminal clones from replicates R6 and R7 against the clones from the initial time point zero stocks confirmed this pattern (Figs. S3 and S4, right panels). The fact that pBR322 has completely reverted to wild-type in R6 (Table S1) and was on the way to purge all variation in R7 suggests that the increase of fitness can be attributed to changes within the bacterial chromosome. Whole genome sequencing of initial and terminal clones from the long term experiments revealed a high frequency nonsynonymous T-to-A transversion in *treB* (Val112Glu) in both replicates (Table S2). In addition, R6 possessed another nonsynonymous substitution in *recA* (Asp161Gly) that effectively re-establishes the RecAi genotype, back to RecA.

The product of the *treB* gene transports trehalose into the cell as trehalose 6-phosphate, which is further converted to glucose 6-phosphate and glucose^42^. In addition to its role as a carbon source, intracellular trehalose can act as an osmoprotectant at high osmolarity ^43–45^. The T-to-A transversion in *treB* observed in our work has been suggested to be a consequence of adaptation of *E. coli* to LB medium and has been identified in three parallel populations in two recent mutation accumulation experiments ^46,47^.

The *recA1*→*recA* reversal in replicate R6 is noteworthy, as it is close to fixation (frequency of 0.91; Table S1) and reinstates recombination ^48–50^. Coincidentally, the same replicate R6 of the long experiment purges plasmid variation completely at the end of sampling, while the other replicate, R7, which did not experience *recA1* → *recA* reversal continues to harbor lingering variation within the plasmid, and has a much lower frequency of the *treB* variant. Recombination has been shown to accelerate adaptation in yeast populations by combining beneficial mutations from different backgrounds and by separating deleterious mutations apart^51^. However, none of the variants we observed are linked, thus it is unclear if RecA has any direct effect on the pattern observed here.

### Modelling the dynamics of plasmid mutations

The observations we described thus far suggest the following scenario. A naïve host is transformed with a plasmid conferring antibiotic resistance. Incubation of the transformed cells in an antibiotic containing environment guarantees retention of an otherwise disposable plasmid. The high density of the turbidostat environment creates a strong selection pressure for shortening generation time.

Initially, selection drives the increase in the frequencies of adaptive changes controlling plasmid copy number, as it translates into the elevated production of the efflux pump, thus helping to offset the inhibitory effect of tetracycline on the protein synthesis. This increases the fitness of cells carrying the mutated pBR322 relative to the wild-type plasmid. However, higher copy number has an associated metabolic cost—once it reaches a certain frequency threshold, it proves too expensive and, in fact, eventually decreases fitness. Simultaneously, a mutation with a large fitness effect arises on the bacterial chromosome. Given the frequencies observed in the experiment, it is highly unlikely that such a chromosomal mutation will arise on the background of an existing plasmid mutations. Its spread to fixation obliterates plasmid variation, as we observed in Fig. 2.

To solidify this reasoning into a quantitative framework, we developed a model drawing on the previous work of De Gelder et al.^52^. Assuming that there is no back mutation on both the plasmid and the chromosome, we denote the number of cells with pBR322 bearing mutations at a certain generation *n* by *m*_*n*_, and those with wild-type pBR322 by *w*_*n*_. At any generation, *m*_*n*_ increases due to (1) doubling of mutant cells from *w*_*n*-1_ occurring at constant rate μ (number of individual cells per generation) and (2) doubling of *m*_*n*-1_with a selection coefficient 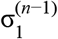. Putting these two components together the number of mutation-carrying cells at generation *n* is given by

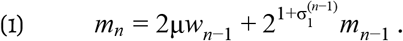

The selection coefficient 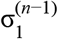 changes from generation to generation due to two factors. First, adaptive plasmid variants increase pBR322 copy number leading to better tetracycline removal ability. This factor is denoted by the constant positive coefficient *b*. Second, increased plasmid copy number imposes a higher burden, inversely proportional to plasmid frequency—characterized as a negative frequency-dependent manner. This effect is denoted by the positive constant *a*. Therefore 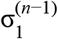 can be expressed as 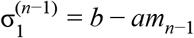. At the same time, the number of cells containing wild-type plasmid *w*_*n*_ increases from previous generation of *w*_*n*−1_ as given by

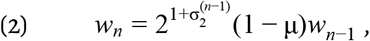

where the selection coefficient 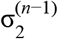 reflects beneficial mutations accumulating on the chromosome and is given by 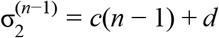 where *c* and *d* are positive constants.

At each sampled time point we can directly observe the fraction of mutants in the population (denoted as β(*n*) as the sum of alternative allele frequencies at sites 3,027-3,035 and 3,118. This corresponds to 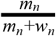, where *m*_*n*_ and *w*_*n*_ are given by equations (1) and (2), and thus β(*n*) is a function of the two selection coefficients 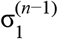 and 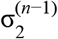, and the mutation rate μ:

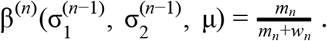

Fitting this model to the empirical data as described in *Material and Methods* helped us explain the observed dynamics of our system. Briefly, we empirically estimated β(*n*) from the observed allele frequencies at each sampling point. We then set biologically plausible intervals of possible values for the *a*, *b*, *c*, *d*, and μ parameters, and employed numerical optimization to infer a set of parameters minimizing the differences between simulated and empirical values of β(*n*) Results of this optimization are given in Table 1. Allele frequencies reconstructed using the described model with optimal parameters closely follow the observed dynamics (Fig. 3) and allow us to make and experimentally test predictions about the behaviour of this system. In particular, it allows for predicting the effects of the increases in tetracycline concentration (Fig. S6) and mutation rate (Fig. S7) on variant frequencies and their trajectories. To model the increase in the tetracycline concentration we decreased the *c* component of the 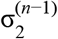 selection coefficient. This effectively increases the beneficial effect of plasmid mutations expressed by 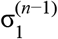 selection coefficient. This increased the maximum frequency of plasmid mutations and prolonged their persistency before ultimately yielding to chromosomal changes. Likewise the effect of increase of the mutation rate allowed for even higher frequencies of plasmid variants. While this dynamics is inherent to our model it provides an insight into the magnitude of selection coefficients and the temporal scale at which plasmid variants remain in the population before being eliminated. This information leads to a series of future experiments our group will be undertaking.

**Table 1.** Optimized estimator for the model.

| Parameter | $a$ | $b$ | $c$ | $d$ | $\mu$ |
| --- | --- | --- | --- | --- | --- |
| Estimator | $1.36 \times 10^{-12}$ | $1.68 \times 10^{-1}$ | $2.69 \times 10^{-3}$ | $3.82 \times 10^{-4}$ | $3.13 \times 10^{-5}$ |

**Figure 3.**
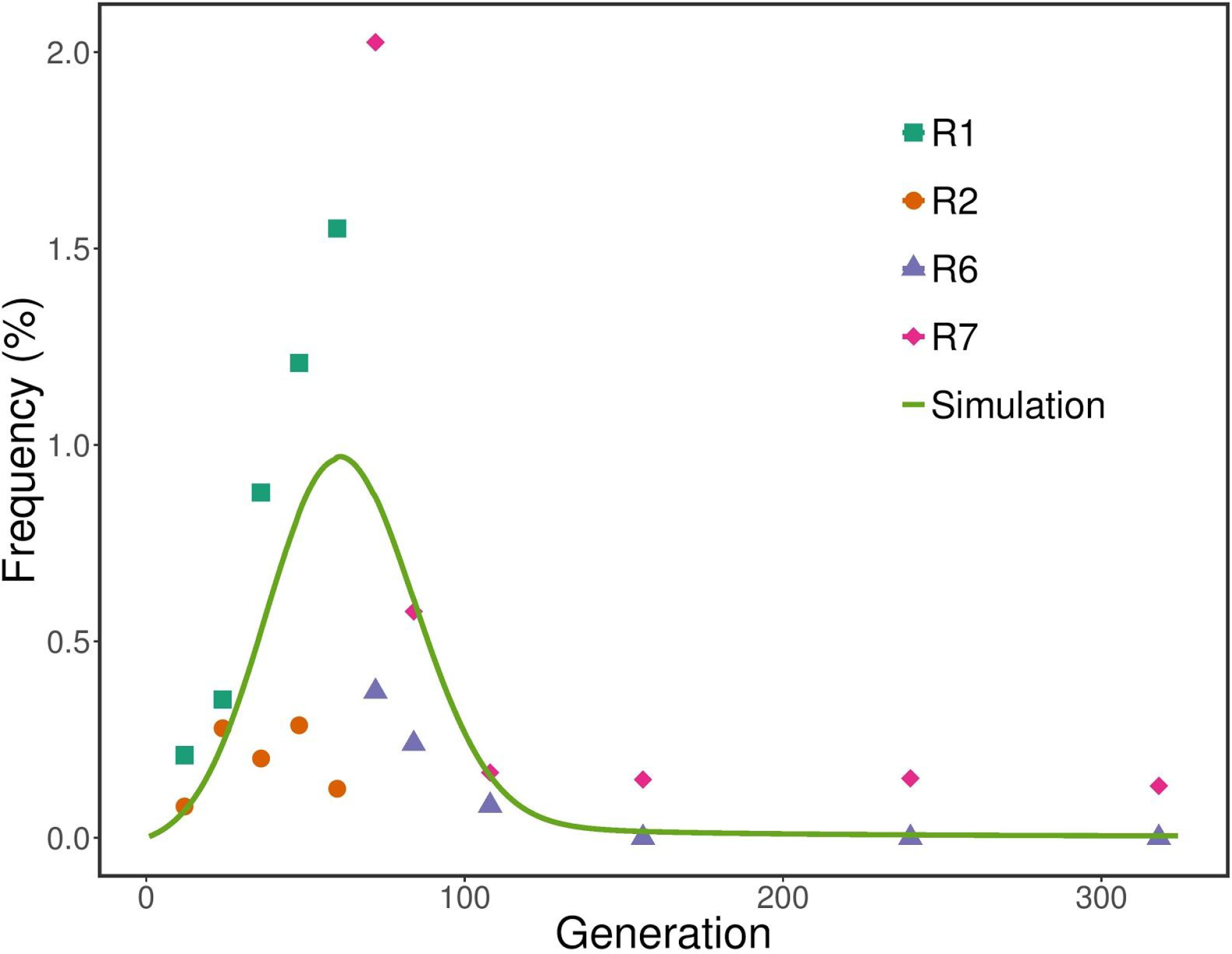
A combination of empirical (R1, R2, R6, and R7) and predicted (simulation) allele frequencies. The simulation was performed with parameters obtained by numerical optimization (see *Material and Methods*). Different turbidostat runs are colored respectively. Y-axis shows the sum of frequencies at bases 3,027-3,035 and 3,118. X-axis represents generations. The x-axis here using generation as the unit corresponds to the x-axis using time in hour in Fig. 1 and Fig. 2. This conversion was accomplished by assuming that the generation time was constantly 60 min.

### Plasmid heteroplasmy likely prevents alterations in copy number

The dynamics described here initially favors beneficial changes within the plasmid DNA that are quickly overridden by chromosomal mutations that attain high frequency in the evolving population. As a result of this co-adaptation process, the plasmid remains unchanged. A number of experimental evolution studies report similar findings where co-adaptation is driven exclusively by chromosomal mutations and leaves plasmids unaltered ^24,53–56^. However, it is not entirely clear why this is the case. At least for pBR322 it is not unreasonable to assume that some combination of mutations affecting plasmid replication could successfully balance tetracycline efflux with copy number burden and reach high frequency at least in a short term. The fact that this does not happen could be a consequence of pBR322 segregation dynamics—the plasmid is partitioned between daughter cells randomly and likely unequally. As pBR322 copy number is only ~20 copies per cell, this sampling error may be significant. Because mutations occur during replication of just one of multiple plasmids copies present within a single cell, this cell is rendered *heteroplasmic*—it contains a mixture of mutated and wild-type pBR322 molecules. The stochasticity of partitioning of plasmid molecules during cell division means that this ratio will be different in each of the daughter cells (also see^57^). In others words they will be heteroplasmic to a different degree. Recent data from another ColE_1_ plasmid (pB_1000_ in *Haemophilus influenzae*) shows heteroplasmic cells to have higher overall number of plasmid copies compared to cells that are monomorphic for the mutated plasmid (here, like in our case, the mutation leads to copy number increase)^58^. Thus the differences in the extent of heteroplasmy (a mutant-to-wild-type ratio within a cell) will lead to different plasmid copy numbers within the daughter cells. This would explain the behavior we observe in our system. Our experiment is performed at a constant antibiotic concentration. This means that the maximum beneficial effects of the increase in the number of efflux pumps is likely associated with a single, “optimal” number of plasmid copies per cell, which balances incurred metabolic cost. It is likely that such optimum is only achieved in heteroplasmic cells because monomorphic mutated cells may produce too many copies of the plasmid to remain competitive in the population. In addition, RNAI, which contains most mutations in our study, acts in *trans* by binding to RNAII at the origin of replication. As a result even a single mutated molecule may influence replication of wild-type molecules within a cell. Thus there must be a narrow optimal value for the degree of heteroplasmy that translates into the optimal pBR322 copy number conferring selective advantage in a given antibiotic concentration. However, the very nature of pBR322 segregation makes it very difficult to reliably establish cells with this “optimal” number in the population because every cell division effectively randomizes the relative numbers of mutated and wild-type plasmids in the next generation. As a result the positive effect of the increase in the copy number is only achieved along a precise optimal value ridge of the adaptive landscape. Every time a cell divides its descendants fall from the ridge because they produce either too few or too many plasmid copies. Likewise ascending the ridge is only possible by chance when a cell division results in just the right degree of heteroplasmy. Because the bacterial populations we are sampling are large, we are able to observe these events until a large fitness effect mutation obliterates the variation by changing the topography of the adaptive landscape (the variation is likely still there but cannot be observed with our method). Addressing these issues will require development of technique allowing tracing individual plasmids in bacterial population (e.g., a finer resolution of the approach recently reported by Rodriguez-Beltran et al.^59^).

In conclusion we have applied the highly sensitive duplex sequencing approach to trace the evolution of pBR322 plasmid during its adaptation to *E. coli* DH5α host. We uncovered previously hidden fine scale dynamics where multiple mutations leading to the increase in the copy number compete ultimately losing to a single chromosomal substitution. Our results suggest that there is a strong incentive not to alter copy number even if it can provide a degree of selective advantage. This incentive is likely rooted in the complex interplay between mutated and wild-type plasmids constrained within a single cell and underscores the importance of understanding of intracellular plasmid variability.

## Material and Methods

### Strains and plasmids

*E. coli* strain DH5α was obtained from Invitrogen (cat. 18265017). The cells were transformed with plasmid pBR322 (NEB cat. N3033S) according to the manufacturer’s protocol. pBR322 carries the replication origin, *tetC*, encoding a tetracycline efflux pump, *bla*, conferring resistance to ampicillin, and *rop*, controlling replication^60^. All experiments were performed in Luria-Bertani (LB) broth (EMD cat. 110285), including those in turbidostat and agar plates, that was supplemented with 30 μg/mL tetracycline (Sigma-Aldrich cat. T_3258_) and 0.05% antifoam B (Sigma-Aldrich cat. A_5757_). Transformants were spread over a LB agar plate. A series of single colonies were picked, grown in LB, and then stored in 20% glycerol stock in a −80°C freezer. One glycerol stock was used as the founder to inoculate all turbidostat runs.

### Turbidostat set-up and experimental evolution

The turbidostat set-up was described by Takahashi et al.^61^. It consists of three connected parts (Fig. S5): a carboy containing LB medium, a bacteria growing chamber that is revolved magnetically, and a waste tank. The carboy and the chamber were pressurized by an aquarium air pump. The volume in the chamber was kept constant at 13 mL. OD_600_ was measured in 30 second intervals and LB was added as necessary to maintain OD_600_ constant at 0.8.

Fifty μL of the glycerol stock was inoculated into 13 mL of LB medium and grown for 14 hours at 250 rpm, 37°C. Four mL cell culture was inoculated into turbidostat. One mL culture was frozen as glycerol stock in −80°C as the initial sample. The remaining culture was labelled as time point 0 and collected by centrifugation at 4,000 g for 5 minutes. The pellet was then transferred to −20°C. Turbidostat runs were carried out in an incubator to maintain 37°C. Eight mL culture was taken from the turbidostat at regular 12 hour intervals, which constitute time points for the subsequent analysis. The turbidostat was immediately refilled by fresh broth to the 13 mL mark. When the turbidostat run was completed, the terminal sample was also collected and archived as glycerol stock in −80°C.

### Population fitness determination

OD_600_ values were constantly monitored, and growth curves were inferred from them. We found that exponential growth between OD_600_ values of 0.6 to 0.8 corresponded to the maximum growth rate, and therefore we used it as a proxy for overall population fitness. Growth rate was defined as

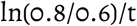

where t is the duration in minutes in which OD_600_ values increased from 0.6 to 0.8. The generation time was calculated as

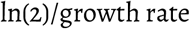

We found that the generation time varied over the course of the two long term replicates. Detailed information regarding growth rate and generation time are displayed in Table S3. For simplicity, the generation time was assumed as 60 min when we modeled the dynamics of variants (Fig. 3).

### Fitness re-measurement of samples representing the initial and terminal time points

Fifty jxL glycerol stock from each of the initial and terminal samples from the two long term replicates R6 and R7 was inoculated into turbidostat in triplicate. OD_600_ was monitored and growth curves were inferred. Exponential growth between OD_600_ values of 0.6 to 0.8 was used to estimate fitness.

### Duplex sequencing and calling plasmid variants

Plasmid at selected time points was extracted by mini-prep (Qiagen cat. 27104). Duplex sequencing library preparation and variant calling were performed as previously described ^20,62^. Briefly, 100 ng of plasmid was sheared to 550 bp on a M220 platform (Covaris) according to the manufacturer’s instructions at the Pennsylvania State University Genomics Core Facility. The sheared DNA was subjected to end-repair (NEB cat. E6050S), size selection (Beckman Coulter cat. A63881), 3′-end dA-tailing (NEB cat. M0212L), duplex adaptor ligation (NEB cat. M0202T), and PCR amplification (Kapa Biosystems cat. kl2612). PCR amplicons were quantified by qPCR (Kapa Biosystems cat. kl4873) and sequenced on an Illumina MiSeq platform using 301-nt paired-end reads. DCS were generated from raw reads and mapped to the reference to call variants. The variant calling workflow is available on Galaxy^63^ at https://usegalaxv.org/u/hanmei/w/du-var-calling.

### Whole-genome sequencing and calling chromosomal variants

Genomic DNA was extracted (Qiagen cat. 51304) from samples representing the initial and terminal time points of the two long term replicates. Whole-genome sequencing was performed as described in our previous article^64^. Briefly, sequencing libraries were prepared as described for duplex libraries, except that 2 μg of DNA was used as the starting material for shearing and adaptors from the Illumina TruSeq Kit were used. Without PCR amplification, ligated libraries were directly subjected to MiSeq sequencing. Variants were called against the DH5α reference genome (GenBank accession number CP017100) using the haploid variant calling workflow on Galaxy at https://usegalaxy.org/u/hanmei/w/hapl0id-var-calling.

### Data modeling

We first defined a *data_generation* function to simulate data based on the mathematical model described by equations (1) and (2) in *Modelling the dynamics of plasmid mutations*. Given the values of the five parameters (*a*, *b*, *c*, *d*, μ), this function returned numbers of mutants *m*_*n*_, number of wild-type cells *w*_*n*_, and mutant frequency, over 318 generations. We also empirically estimated the mutant frequency 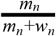 at the generations corresponding to the time points selected for plasmid duplex sequencing. In particular, at each of these empirically determined generations, we computed the empirical mutant frequency as the sum of frequencies of mutations in bases 3,027-3,035 and 3,118 determined in turbidostat experiments. We then defined the *loss* function as the sum of squared differences between simulated and empirical mutation frequencies.

This *loss* function was optimized over the five parameters (*a*, *b*, *c*, *d*, μ) using the R function *optim*, setting the bounds on the parameters as shown in Table S4 (choice of these bound is discussed in the next paragraph). Note that *optim* was run with a maximum of 10^6^ iterations in order to facilitate convergence of the optimization process. Since the simulated mutant frequency shows a complex dependence on the parameters (*a*, *b*, *c*, *d*, μ), the *loss* function is likely to have multiple local minima. As a consequence, the set of optimal parameters (*a*, *b*, *c*, *d*, μ) returned by of the *optim* function strongly depend on their initial values. To effectively explore the parameter space, we randomly sampled initial values for the five parameters (uniformly on a log_10_ scale, according to the bounds in Table S2) and repeated the optimization process until 1,000 converged optimization processes were obtained. Each repetition returned a set of parameters and the corresponding *loss* value. The five values (*a*, *b*, *c*, *d*, μ) generating the smallest *loss* among the 1,000 obtained were chosen as optimal parameters and presented in Table 1. The steps implementing parameters optimization are described in the supplementary jupyter notebook “Figure_3_parameter_optimization.ipynb” at https://github.com/hanmeiqi9l/pBR322-variant-dvnamics.

We defined the five parameter intervals in logarithmic scale to ensure that they were wide enough to randomly sample the parameters. Therefore, a large parameter space was explored. The mutation rate μ in De Gelder et al.^52^ was determined with the order of magnitude of 10^−5^. We set the interval for μ from 10^−10^ to 10^−2^. The other four parameters (*a*, *b*, *c*, *d*) were involved in calculation of 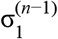 and 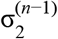. As coefficients, 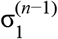 and 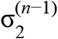 were generally expected to be smaller than 1. We set the intervals for (*b*, *c*, *d*) the same as μ. The interval for parameter *a* was set on smaller values (from 10^−14^ to 10^−6^) because it served as the coefficient of *m*_*n*-1_ and we tried to maintain the product *am*_*n*-1_ reasonably small.

## Data deposition

Raw sequencing reads in this study have been deposited at the NCBI SRA database as BioProject PRJNA485503.

## Supplementary files

Supplementary files include Table S1 (xlsx), Table S2 (xlsx), Table S3 (xlsx), and Table S4 (xlsx), jupyter notebooks and data to perform the parameter optimization shown in Fig. 3 and to generate predictions shown in Figs. S6 and S7. The notebooks can be found at https://github.com/hanmeiqi9l/pBR322-variant-dvnamics.

## Acknowledgements

Authors are grateful to Jim Bull for his suggestions that have greatly improved our manuscript. Nick Stoler has aided in the tuning of the duplex data analysis pipeline. This study has been funded by the funds provided by the Eberly College of Science at the Pennsylvania State University and NIH Grants U41 HG006620 and R01AI134384-01 as well as NSF ABI Grant 1661497.

---

**Supplementary Figure 1.**
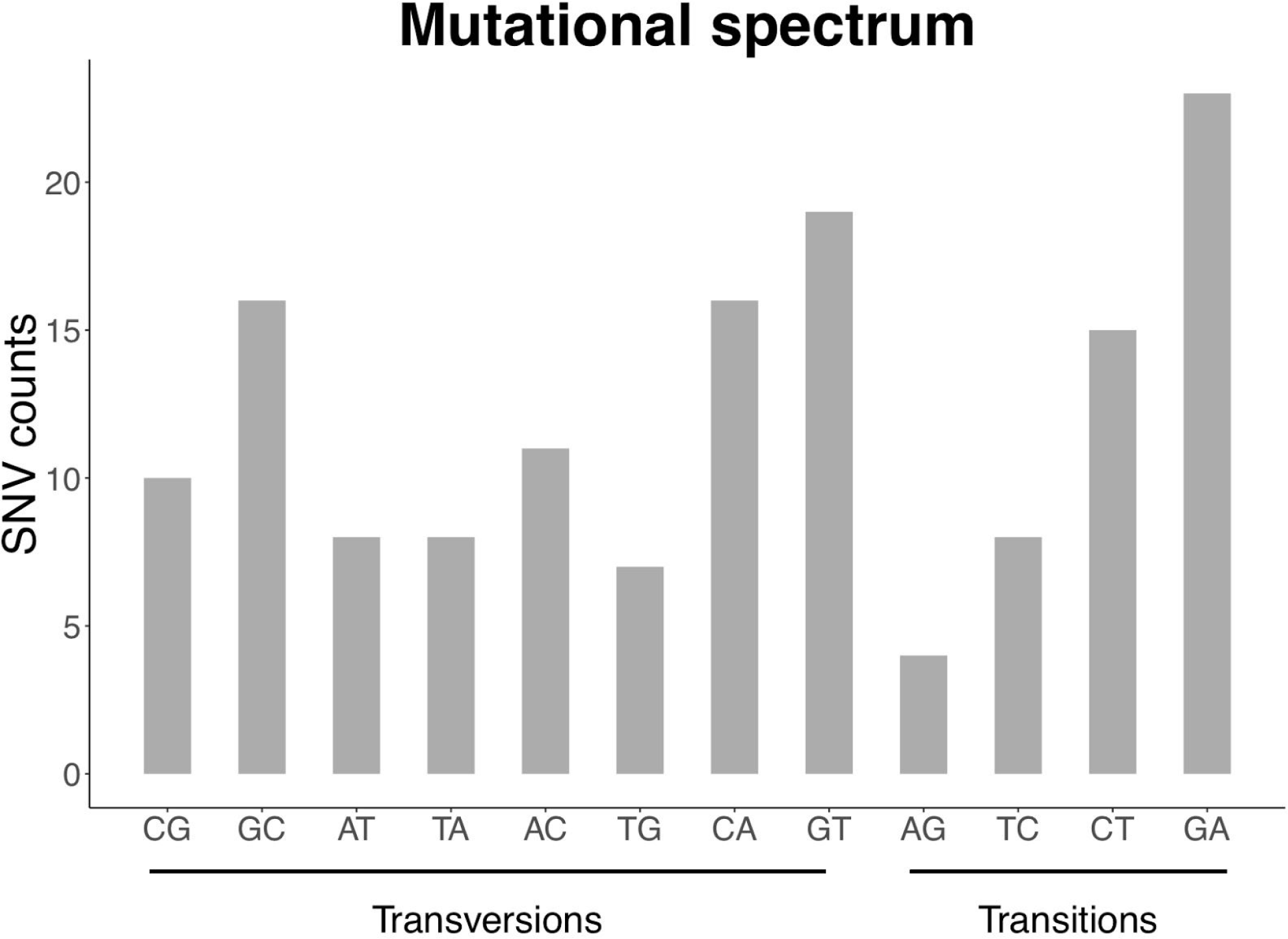
Mutational spectrum across 145 SNVs relative to the reference sequence of pBR322 in our experiment. X-axis shows different mutation types. Y-axis shows the number of SNVs falling into respective mutation types.

**Supplementary Figure 2.**
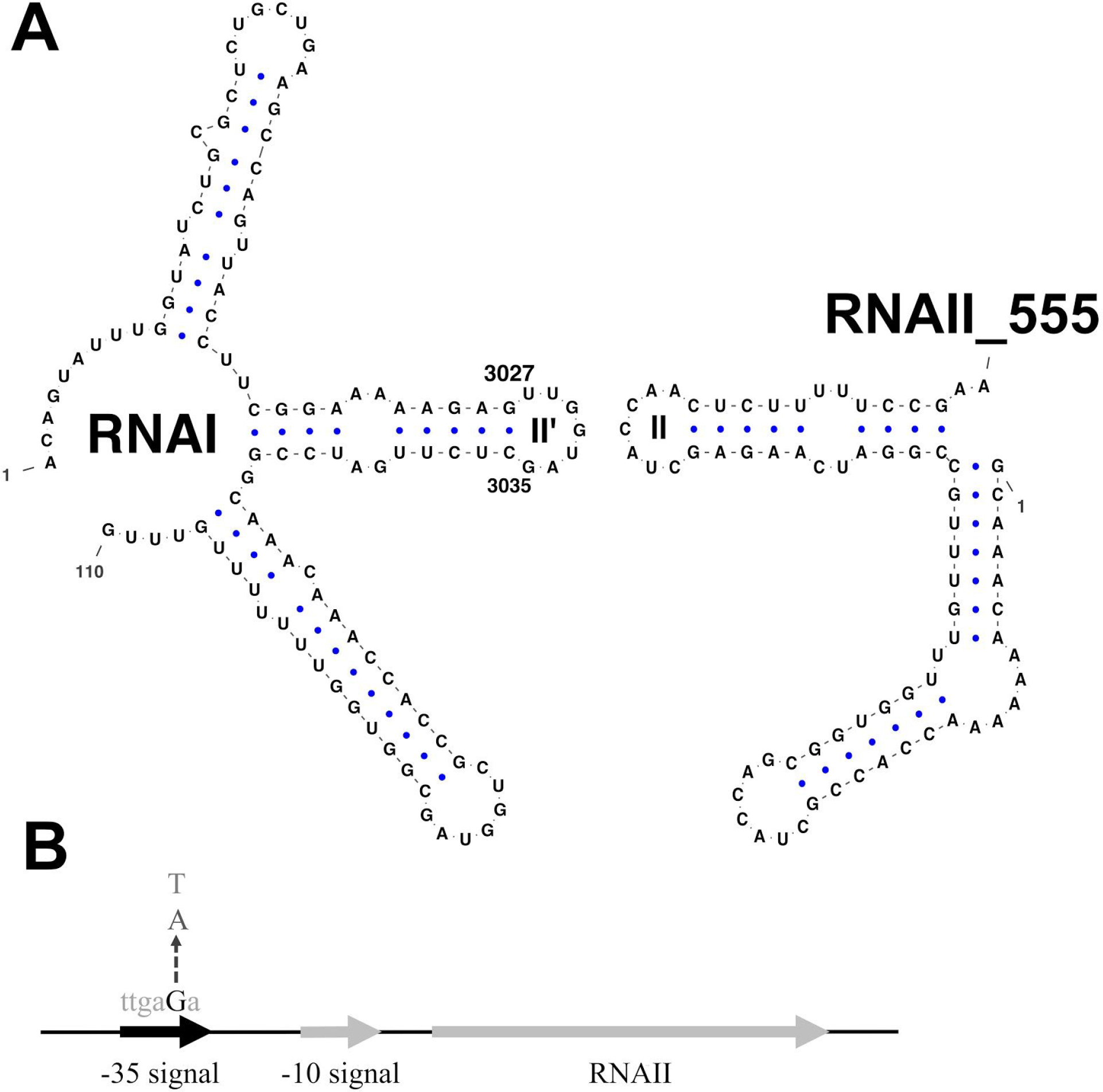
Secondary structures of RNAI and RNAII molecules. **A**. Positions 3,027 and 3,035 are boundaries of the region that contains the majority of beneficial mutations identified in this experiment. RNAII was reported 555 nt in length. Only RNAII loop regions interacting with RNAI are shown, and denoted as “RNAII_555”. **B**. Mutations at base 3,118 (uppercase, black) increase transcription of RNAII.

**Supplementary Figure 3.**
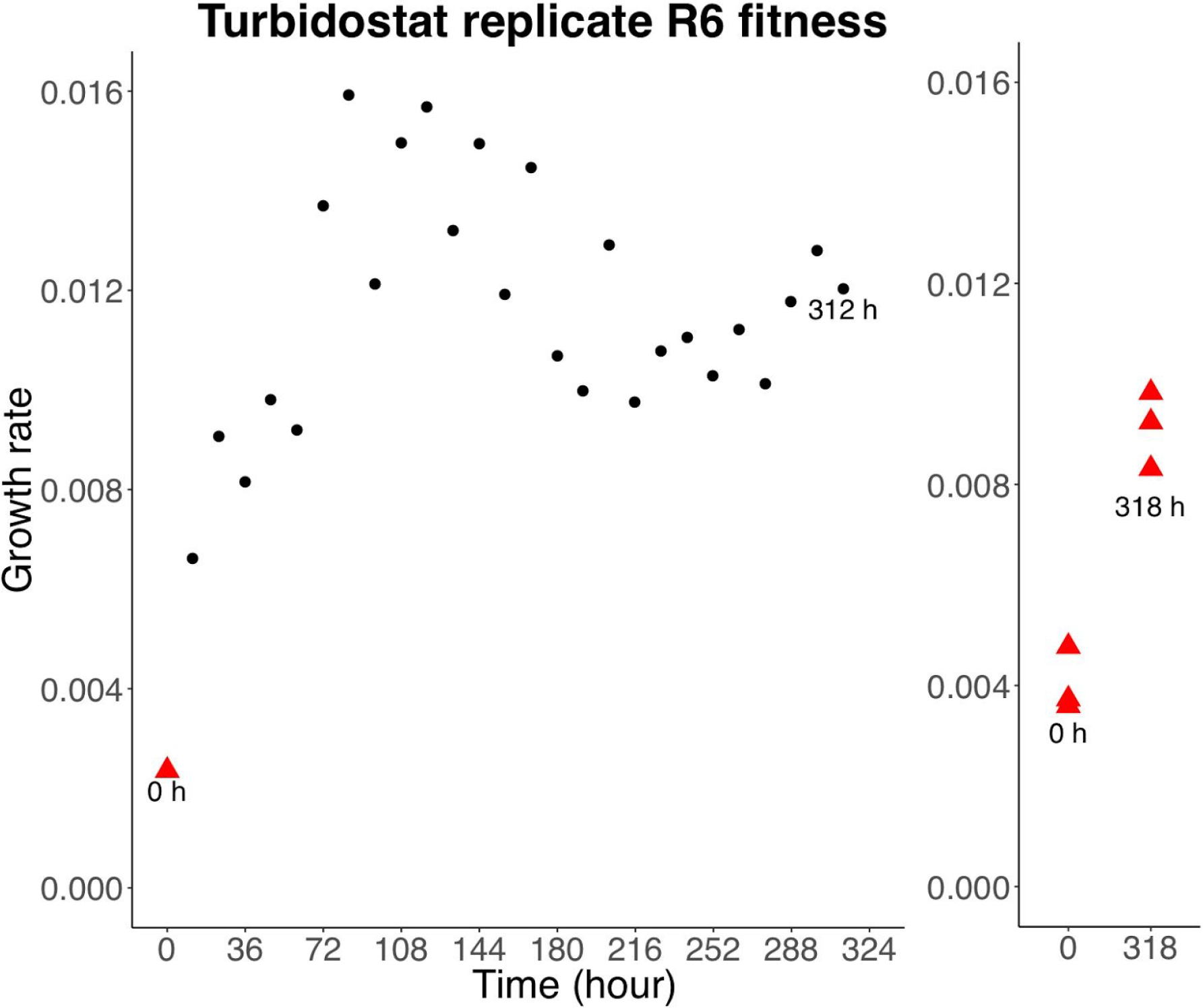
Growth rate of individual isolates from replicate R6 of the long turbidostat run. In the left panel, growth rate were estimated from slopes in Fig. 1. Growth rate the initial isolate is shown in a red triangle, and those of the remaining isolated are shown in black circles. Growth rate of the terminal isolate in replicate R6 cannot be obtained from Fig. 1, because the terminal isolate was stored in the glycerol stock and did not show a dip in OD values. However, it can be re-measured by inoculating the glycerol stock into turbidostat; this is shown in the right panel. In the right panel, initial and terminal isolated, both in red triangles, were re-measured in triplicates.

**Supplementary Figure 4.**
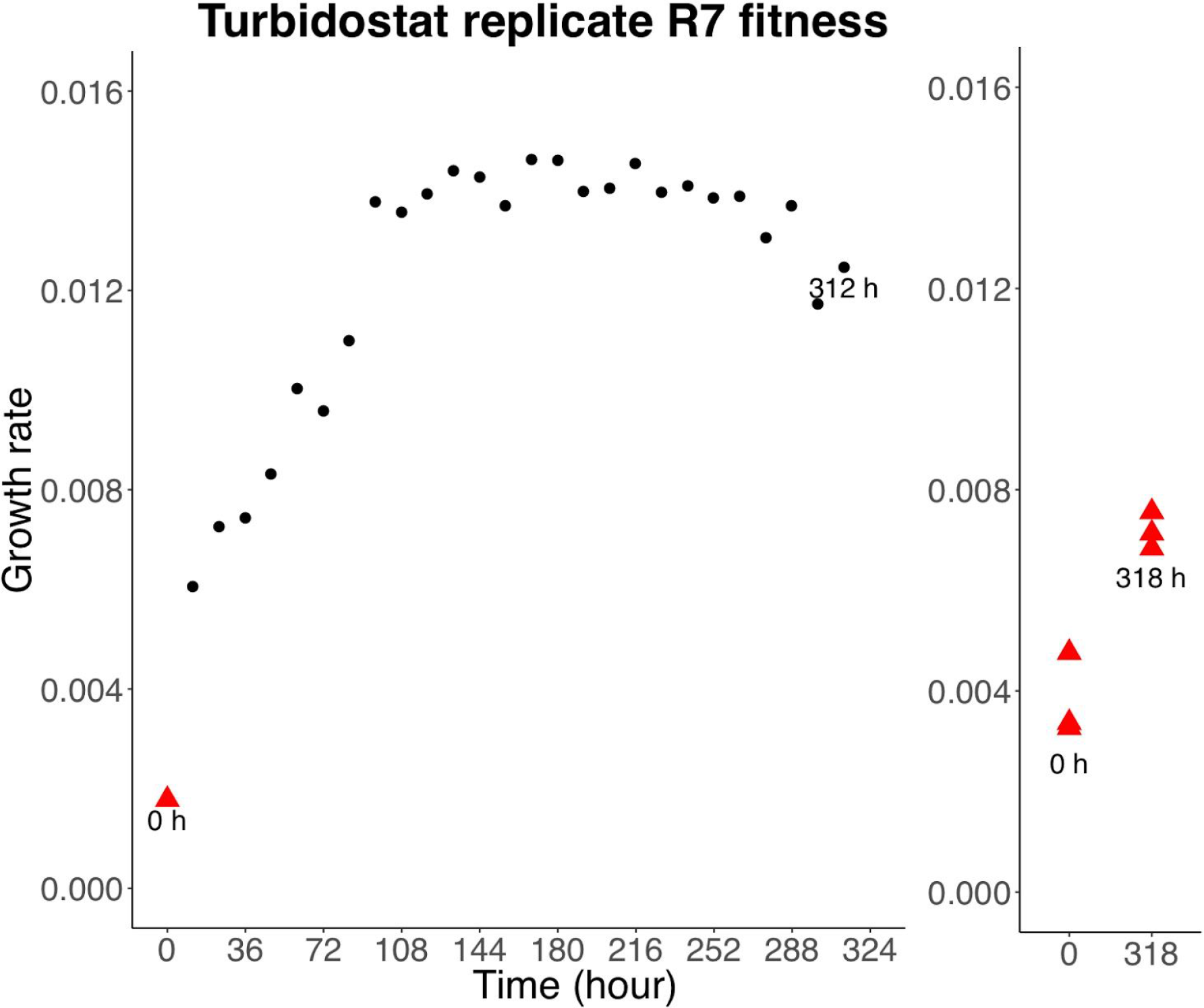
Same as Fig. S3, growth rate of individual isolates from replicate R7 of the long turbidostat run.

**Supplementary Figure 5.**
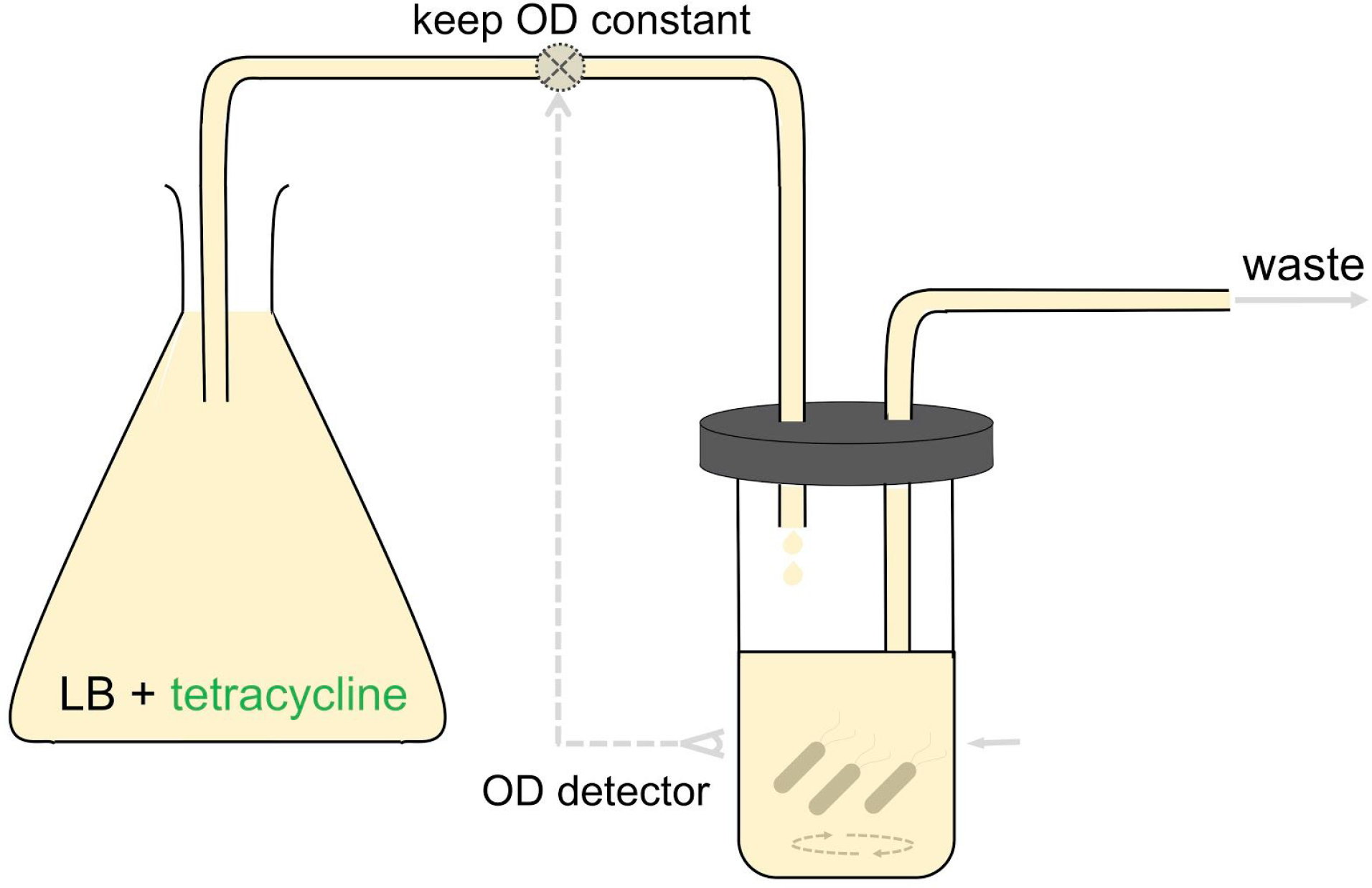
Turbidostat setup.

**Supplementary Figure 6.**
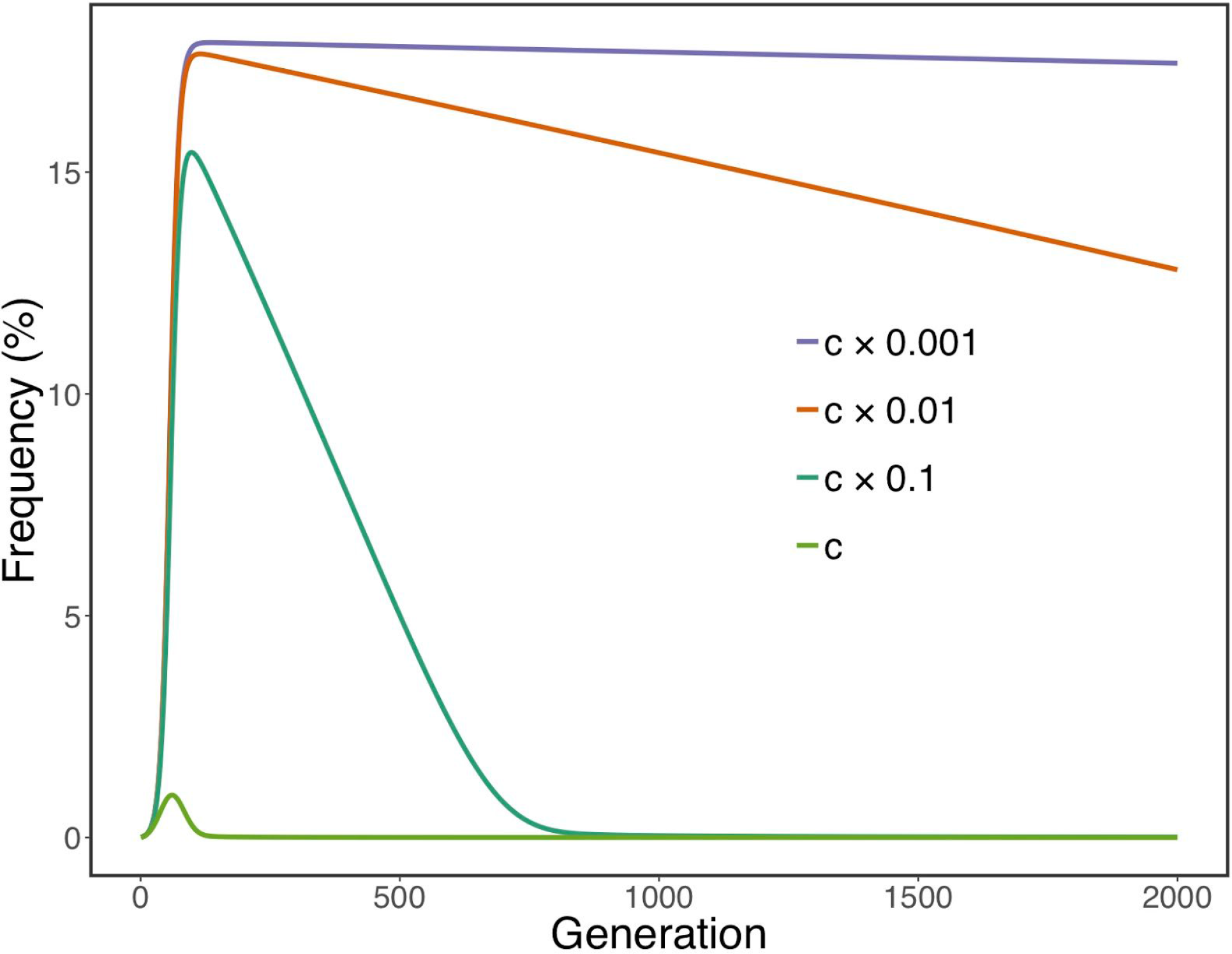
Prediction of frequencies of lineages carrying plasmid mutations at bases 3,27-3,035 and 3,118. The *a*, *b*, *d*, μ parameters were given the same value as those obtained by numerical optimization and used in Fig. 3. The *c* parameter was multiplied by 0.1, 0.01, and 0.001. Different *c* values generated different dynamics, which were shown in different colors.

**Supplementary Figure 7.**
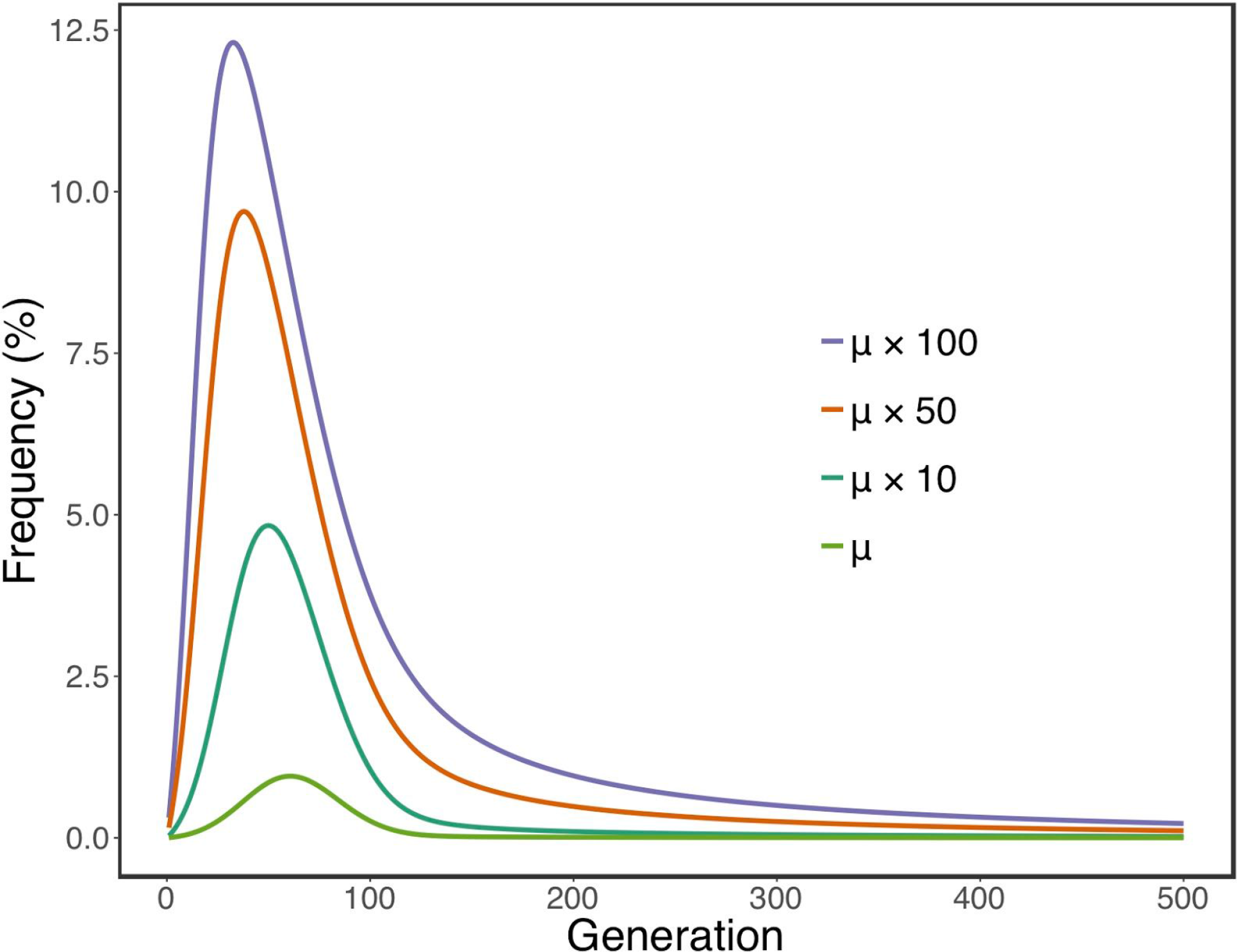
Same as Fig. S6. The *a*, *b*, *c*, *d* parameters were given the same value as those obtained by numerical optimization. The (x parameter was multiplied by 10,50,and 100, generating different dynamics.

**Table S1.** Plasmid SNVs and INDELs identified in all four turbidostat runs.

**Table S2.** Chromosomal mutations identified in the two long term replicatesR6 and R7.

**Table S3.** Growth rate and generation time over the course of the two long term turbidostat replicates.

**Table S4.** Bounds, in log_10_ scale, for the parameters to be drawn uniformly at random to model the dynamics of plasmid.

